# Effect of prolonged captivity on the hemolymph profile of *Tachypleus gigas*

**DOI:** 10.1101/2020.04.24.059816

**Authors:** B. Akbar John, Hassan I Sheikh, J.A. Ichwan Solachuddin, B.Y. Kamaruzzaaman

**Affiliations:** Institute of Oceanography and Maritime Studies (INOCEM), Kulliyyah of Science, International Islamic University Malaysia (IIUM), Kuantan, Pahang Malaysia; Faculty of Fisheries and Food Science, Universiti Malaysia Terengganu, 21030, Kuala Nerus, Terengganu, Malaysia

**Keywords:** *Tachypleus gigas*, anticoagulant, living fossil, amebocyte cells, captive breeding

## Abstract

Horseshoe crabs amebocyte cells degranulate to form a gel clot when in contact with endotoxins. This phenomenon is the basis of both Horseshoe crab immune system and detection of endotoxin in biologicals. The present study investigates the amebocyte cells quality in *Tachypleus gigas* pre and post bleeding under captivity. Wild and captive horseshoe crabs (5 months captivity) were bled in 6 anticoagulant formulations (A, B, C, D, E and F). No profound difference in cell density between captive and wild groups with the mean value of 0.883×10^7^ a cells/mL and 0.917×10^7^ cells/mL, respectively. while, the cell viability of the captive group was significantly lower than the wild crabs (*F*=808.075, p<0.001). Anticoagulant formulation significantly affected cell viability and cell morphology in both captive and wild groups (*p*<0.001). Amebocyte cells collected from the wild *T. gigas* using optimum anticoagulant (formula C) showed 0.6 ×10^7^ cells/mL cell density and 86.9% cell viability, while morphology analysis revealed the percentage of contracted, granular flattened and degranulated flattened cells were 14.62%, 71.39% and 14%, respectively. The anticoagulant formulations showed varying capabilities in maintaining cell viability due to its buffering and chelating capacity. We conclude that captivity has a negative effect on the amebocyte cell quality.

**HIGHLIGHTS:**

- Captivity has a negative impact on amebocyte quality in horseshoe crab (*Tachypleus gigas*).
- No significant difference in cell density between captive and wild crabs.
- Anticoagulant formulations have significant impact on the cell density, viability and morphology
- Captive crabs were immunocompromised due to single specific feed intake.

## INTRODUCTION

Horseshoe crab has been exploited by several industries which raised genuine concerns about the future of this living fossil. The biggest threats are habitat destruction and uncontrolled harvesting for fisheries and biomedical industries (Eyler, 2016; John et al., 2018a; John et al., 2018b; Krisfalusi-Gannon et al., 2018). Pharmaceutical companies produce Limulus/Tachypleus Amebocyte Lysate (LAL/TAL) from the blood of horseshoe crab which is used to detect live-threatening endotoxins (Burger & Gochfeld, 2000; Kreamer & Michels, 2009). However, bleeding horseshoe crab for LAL production has caused mortality rate as high as 30% (Kumar, Roy, Sahoo, Behera, & Sharma, 2015; Novitsky, 2015). TAL production led to 100% mortality of as many as 600,000 crabs that were harvested in China, bled to death, then sold in restaurants for human consumption (Gauvry, 2015). These facts could explain the decline in populations and indicates unsustainable fishery practice of horseshoe crabs for the pharmaceutical industry especially in Asia.

Though it is hard to predict the exact market value of the LAL/TAL industry in the near future, the size of the pharmaceutical market it serves is exceeding $1.2 trillion in the year 2018 with an average yearly increase of ~5.8%. Global vaccine sale is projected to increase over 50% to $100 billion USD by 2025 from the current sale of $49 billion USD in 2018 (Cramer, 2015; WHO, 2019). This increased utilization is expected to increase the pressure on the horseshoe crab populations. Alternative options to bleeding practices includes the recently approval recombinant Factor C (rFC) by Food and Drug Administration (FDA) (Maloney, Phelan, & Simmons, 2018). Aquaculture and captive breeding practices is another alternative that could aid in the conservation of horseshoe crab. In this approach, it is proposed that a biomedical company can have a limited and predetermined number of horseshoe crabs for LAL/TAL industry. This approach not only protects the wild horseshoe crab populations, but also eliminates the quantitative as well as qualitative variations reported between various manufacturers and batches (Hurton, Berkson, & Smith, 2005; Novitsky, 2009). However, captivity has been reported to have various effects on horseshoe crabs that indicate health decline and can be detrimental to the quality of amebocyte lysate produced from captive horseshoe crab (Coates, Bradford, Krome, & Nairn, 2012; Prior, 1990).

During the bleeding practice, spontaneous degranulation of amebocyte cells, exocytosis and clot formation are unavoidable and can significantly impact haemolymph quality. In order to minimize coagulation, numerous chemicals and formulations have been proposed and used as an anticoagulant. For example, caffeine (Nakamura, Morita, & Iwanaga, 1985), propranolol (Cohen et al., 1985), dimethylsulfoxide (Liang & Liu, 1982), sulfhydryl reagents (Cohen et al., 1985), membrane-active detergent Tween-20 (Nicetto et al., 2019; Strickley et al., 2019) in different concentrations of LPS-free NaCl solution are commonly used to mimic the blood plasma condition.

In the case of horseshoe crab, numerous anticoagulants have been reported with varying pH, buffering capacities and ingredients (Armstrong & Conrad, 2008). Therefore, several anticoagulant formulations were selected from literature and modified in this study. Various aspects of ideal anticoagulant were examined such as chelation capacity, buffering capacity, pH, osmotic pressure regulation and energy supply. Citric acid, EDTA (Ethylenediaminetetraacetic acid) and disodium ethylenediaminetetraacetic acid (Na2EDTA) were used in order to chelate free calcium ions that trigger coagulation and stabilize the cell (Acton, 2013; Li, Zhang, & Francis, 2013; Ratcliffe, 1993). The pH levels selected were 4.6, 7.5 and 8.77. The importance of energy supply was also examined by selectively providing dextrose as a carbon source to supply amebocyte cells with adequate energy to maintain its cellular activities (Li et al., 2013).

In order to evaluate the effect of captivity, published literatures reported that pharmaceutical companies bleed horseshoe crabs only twice a year, hence 5-6 months is considered to be a sufficient time for the horseshoe crab to restore its blood count (Cramer, 2015). In captivity, Armstrong and Conrad (2008) suggested that horseshoe crabs can be bled up to twice a month. However, earlier studies indicated that horseshoe crabs that are kept in captivity need 6 months to restore amebocyte cells count (Prior, 1990). Hence, in this study, the horseshoe crabs were captured, transported and bled on the sampling date, therefore, mimicking current bleeding practices employed by biomedical companies. This group was labelled as “wild group” and is considered to have the ideal quantity and quality of amebocyte cells. Then, the bled horseshoe crabs were kept in captivity for 5 months, as suggested by literature to be sufficient time to restore their amebocyte cell count and quality (Armstrong & Conrad, 2008; Cramer, 2015).

Aquaculture techniques guarantee continuous food supply as well as control over crucial parameters such as pH, temperature and salinity which were all maintained at similar levels to that of the sampling area (Balok beach). With continuous feeding, clean water and stable environmental parameters, horseshoe crabs were assumed to be in ideal health. Experimental evidence has shown profound negative effects of captivity on the hemolymph quality of juvenile *Tachypleus tridentatus* (Kwan, Chan, Cheung, & Shin, 2014).

Hence, the present study will examine the effect of captivity (5 months) on the quality of horseshoe crabs’ amebocyte cells and address the effect of different anticoagulants formulations on the hemolymph quality.

## EXPERIMENTAL DESIGN

### Horseshoe crab samples

Horseshoe crab (*Tachypleus gigas*) were collected from Balok beach (Latitude 3°58.194” N, Longitude 103°22.608” E) Malaysia. The average sea parameters in Balok beach were 26.13 °C ± 2.23 temperature, 7.57 ± 0.65 pH and 24.88 ± 4.12 ppt salinity throughout non-monsoon. Ambient water parameters on sampling day were 27.6 °C temperature, 8.35 pH and 24.05 ppt salinity. A total of 18 female horseshoe crabs (carapace width 21.92±1.17cm) were collected using gill net and transported to the laboratory and acclimated for a day before bleeding. Crabs were tagged with different coloured Nylon66 cable zip tie tags and morphometric measures (Total Length, Carapace Width, Interocular Width, Tail Length, Body Weight) were documented (Supplementary file 1). Three crabs for each anticoagulant treatment were bled immediately and were considered as “wild group”.

### Captive condition

Eighteen *T. gigas* were then maintained in captivity for 5 months at the INOCEM Research Station (IRS), Pahang, Malaysia, and were considered as “captive group”. All tanks were maintained at 26 ± 1 °C temperature, 7.5 ± 0.2 pH, salinity of 30 ± 2 ppt, dissolved oxygen: 6.0±1 mg/L and ammonia concentration below 0.05 mg/L. Horseshoe crabs were fed 3% of their body weight with bivalves (*Meretrix meretrix*) which was equivalent to 2 clams/crab/day. The proximate composition of feed was measured by ProxiMate™ (Protein: 13.17-15.65%; Lipid: 1.21-1.78%; Ash: 2.87-3.62%; Carbohydrate: 5.11-6.37% and Moisture: 76-79.31%). The aquarium was filled with coarse sand to mimic the natural environment of the horseshoe crab. The sand was cleaned/replaced every 2-3 days and water exchanged daily. The aquarium diameters were 92” × 48” × 30” (L × W × D). The density of the crabs in the aquarium was 2 pairs/square meter.

### Bleeding regime

All salts and glassware used were depyrogenated at 180 °C for 4 hrs. The arthroidal membrane was wiped with 70% ethanol using Kim-wipes before and after hemolymph extraction. Hemolymph withdrawal was carried out using a pre-chilled 18-gauge needle (Terumo) inserted into the cardiac sinus of the horseshoe crabs following strict sterility measures. The hemolymph was collected in falcon tubes containing various pre-chilled marine anti-coagulant formulations (1:1, Blood: Anticoagulant).

### Anticoagulants formulations

Formula **A** [adopted and modified from John (2012)]:

> 9 mM Disodium Ethylenediaminetetraacetic Acid (Na_2_EDTA, R&M Chemicals), 0.75 mM Sodium Chloride (NaCl, QrecTM), 0.1 M Dextrose (Merck), pH 4.6.

Formula **B** [adopted and modified from Coates, Whalley, and Nairn (2012)]:

> 0.51 M Sodium Chloride (NaCl), 47 mM Citric Acid (Sigma) and 10 mM Disodium Ethylenediamine-tetraacetic Acid (Na_2_EDTA), 0.1 M Dextrose, pH 4.6.

Formula [adopted from Armstrong and Conrad (2008)]:

> 30 mM Sodium Citrate (R&M Chemicals), 26 mM Citric Acid, 10 mM Disodium Ethylenediamine-tetraacetic Acid (Na_2_EDTA), 0.1 M Dextrose, pH 4.6.

Formula **D** [adopted and modified from (Armstrong & Conrad, 2008; Liang & Liu, 1982)]:

> 1 mM Na_2_EDTA, 70 mM DMSO (Sigma-Aldrich), pH 4.6.

Formula **E** [modified from HEPES Good’s buffer]:

> 0.1 M HEPES (Sigma-Aldrich), 10mM Na_2_EDTA, 0.1 M Dextrose, pH 7.4.

Formula **F** [modified Bicarbonate-Carbonate buffer]:

> 0.1 M Na_2_CO_3_ (R&M Chemicals), 0.1M NaHCO_3_ (R&M Chemicals), 10 mM Na_2_EDTA, 0.1 M Dextrose, pH 8.77 (R&M)

### Evaluation of amebocyte cells quality

Viability of amebocyte cells determined in all six anticoagulant formulations using trypan blue dye (Sigma-Aldrich) similar to Coates, Bradford, et al. (2012). amebocyte cells were diluted in the same anticoagulant formulation used in bleeding. An equal volume (500μl) of 0.4 trypan blue was added to the amebocyte cells and were allowed to mix for 30 minutes. amebocyte counts were estimated in triplicate using an improved Neubauer hemocytometer (Labour optics). Stained (dead) vs unstained (alive) were used to calculate the percentage of viability. Nikon ECLIPSE DS-Ri1 microscope (NIS Element imaging software V 4.13) was used to observe and count amebocyte cells in brightfield settings at 40x magnification. Cell morphology was analysed by diluting hemolymph (500μl) in an equal volume of pre-chilled anticoagulants supplemented with 2.5% neutral formaldehyde. Serial dilutions were carried out in LPS-free saline (3% NaCl, 10 mM NaHCO3, pH 7.5). Morphology of 30-50 cells were counted in each replicate and multiplied with dilution factor in order to ensure data reliability. The cells were viewed under a microscope using bright-field settings at 40x magnification. The cells’ morphology were labelled following Hurton et al. (2005) classification as contracted (C), granular-flattened (GF) and de-granular-flattened (DF) (Figure 1).

**Figure 1.**
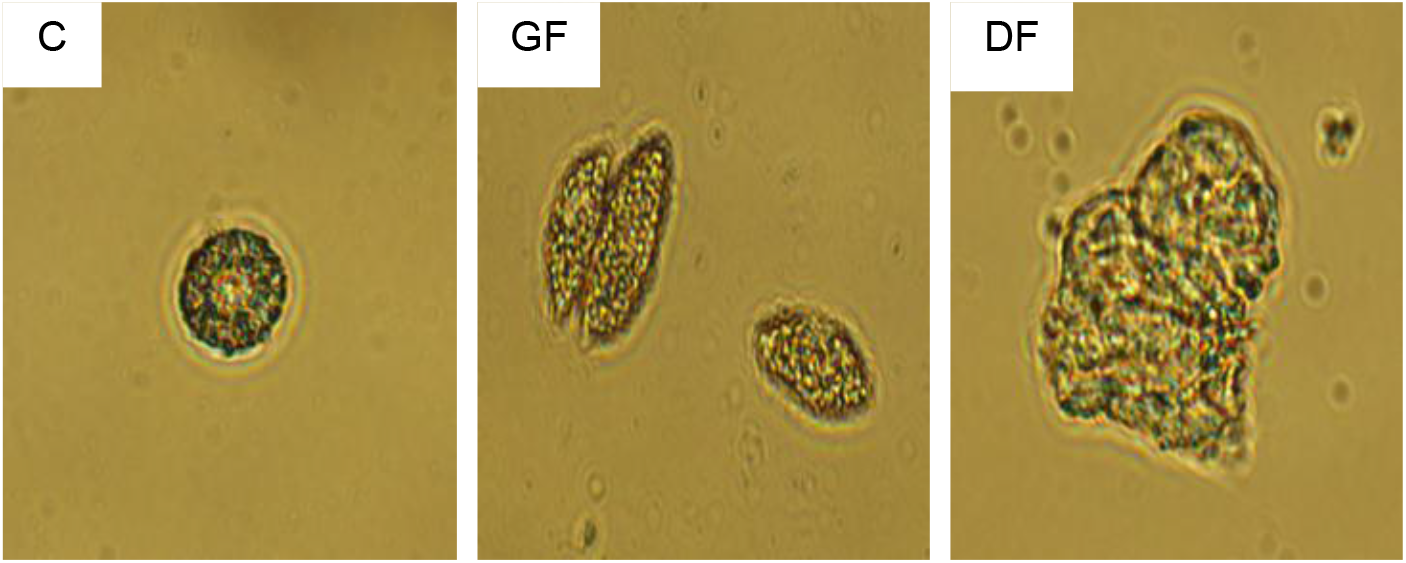
Characterization of amoebocyte cells shapes observed in this study. C: Contracted, GF: Granular flattened, DGF: Degranulated flattened.

### Statistical analysis

Hemolymph quality (amoebocyte cell density, viability and morphology) of wild and captive crabs obtained in various anticoagulants (A, B, C, D, E, F) were analysed using SPSS v19. The effect of captivity and difference between anticoagulant formulations were compared using general linear model and Tukey’s post-hoc analysis. This is due to independent factor having multiple subcategories (captivity and anticoagulant) and comparison of several dependent variables (amoebocyte cells’ density, viability and morphology). Amoebocyte cells’ morphology was divided into contracted (C), granular flattened (GF) and degranulated flattened (DF) which were measured in percentage. Data were expressed in mean ± standard deviation (SD). Statistical significance was expressed at 95% confidence level (p<0.05), 99% confidence level (p < 0.01) or 99.9% confidence level (p < 0.001).

## RESULTS AND DISCUSSION

### Effect of captivity on horseshoe crab hemolymph

Table 1 summarizes the hemolymph quality of captive and wild *T. gigas* obtained in this study. Two independent studies have reported that the amebocyte cell density in a healthy *Limulus polyphemus* was between 2-6×10^7^ cells/mL (Coates, Bradford, et al., 2012; Levin & Bang, 1968). However, in the present study, the mean amebocyte cell density in wild *T. gigas* was 0.883×10^7^ cells/mL. This can be due to age and/or size of *T. gigas* compared to *Limulus polyphemus*. For instance, amoebocyte cell density of of wild juvenile *T. tridentatus* and *C. rotundicauda* (8^th^ to 11^th^ instar) ranged from 0.26 to 0.53 ×10^7^ and 0.14 to 0.43 ×10^7^ cells/mL, respectively (Kwan et al., 2014).

**Table 1:**
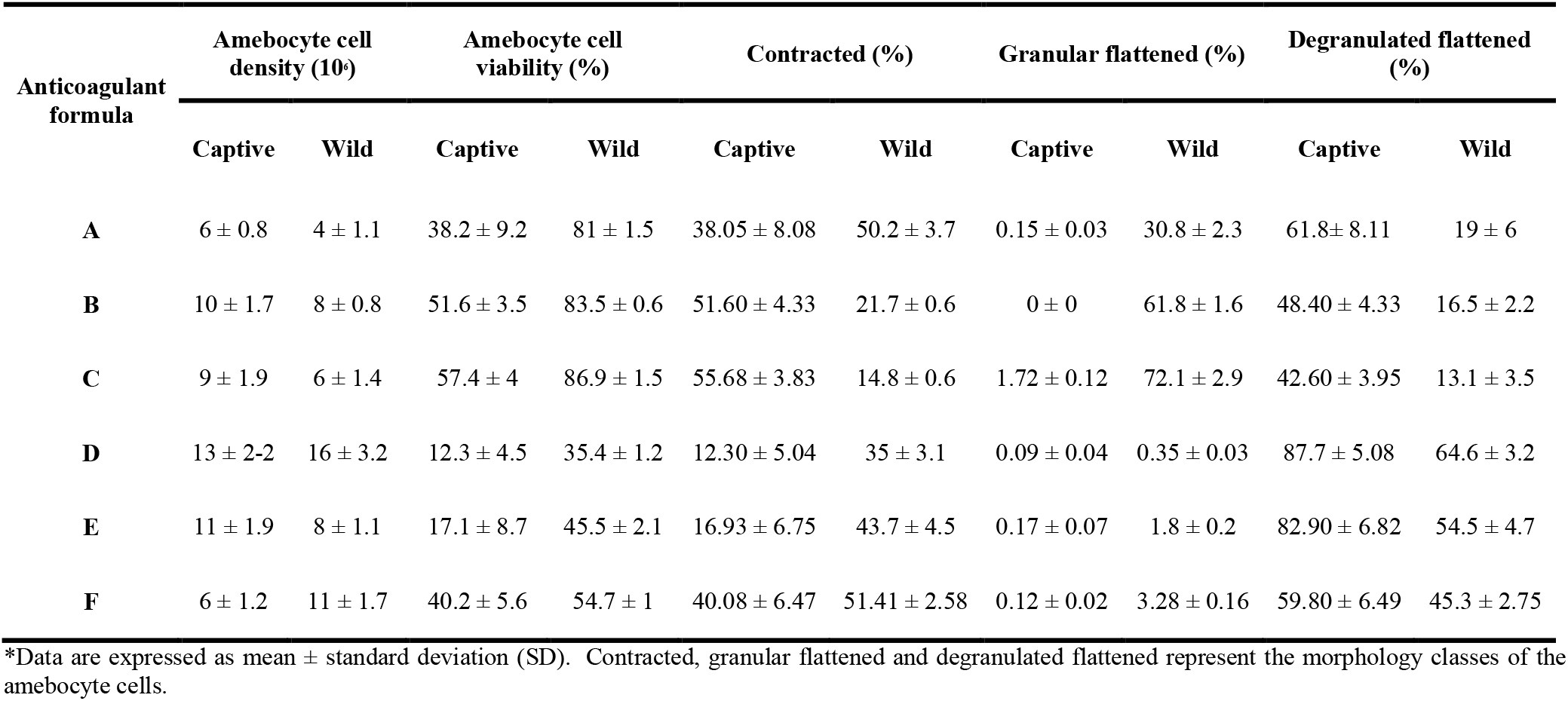
Hemolymph quality of captive and wild *Tachypleus gigas*

| Anticoagulant<br>formula | Amebocyte cell<br>density (10<sup>6</sup>) |  | Amebocyte cell<br>viability (%) |  | Contracted (%) |  | Granular flattened (%) |  | Degranulated flattened<br>(%) |  |
| --- | --- | --- | --- | --- | --- | --- | --- | --- | --- | --- |
|  | Captive | Wild | Captive | Wild | Captive | Wild | Captive | Wild | Captive | Wild |
| **A** | 6 ± 0.8 | 4 ± 1.1 | 38.2 ± 9.2 | 81 ± 1.5 | 38.05 ± 8.08 | 50.2 ± 3.7 | 0.15 ± 0.03 | 30.8 ± 2.3 | 61.8 ± 8.11 | 19 ± 6 |
| **B** | 10 ± 1.7 | 8 ± 0.8 | 51.6 ± 3.5 | 83.5 ± 0.6 | 51.60 ± 4.33 | 21.7 ± 0.6 | 0 ± 0 | 61.8 ± 1.6 | 48.40 ± 4.33 | 16.5 ± 2.2 |
| **C** | 9 ± 1.9 | 6 ± 1.4 | 57.4 ± 4 | 86.9 ± 1.5 | 55.68 ± 3.83 | 14.8 ± 0.6 | 1.72 ± 0.12 | 72.1 ± 2.9 | 42.60 ± 3.95 | 13.1 ± 3.5 |
| **D** | 13 ± 2-2 | 16 ± 3.2 | 12.3 ± 4.5 | 35.4 ± 1.2 | 12.30 ± 5.04 | 35 ± 3.1 | 0.09 ± 0.04 | 0.35 ± 0.03 | 87.7 ± 5.08 | 64.6 ± 3.2 |
| **E** | 11 ± 1.9 | 8 ± 1.1 | 17.1 ± 8.7 | 45.5 ± 2.1 | 16.93 ± 6.75 | 43.7 ± 4.5 | 0.17 ± 0.07 | 1.8 ± 0.2 | 82.90 ± 6.82 | 54.5 ± 4.7 |
| **F** | 6 ± 1.2 | 11 ± 1.7 | 40.2 ± 5.6 | 54.7 ± 1 | 40.08 ± 6.47 | 51.41 ± 2.58 | 0.12 ± 0.02 | 3.28 ± 0.16 | 59.80 ± 6.49 | 45.3 ± 2.75 |

\*Data are expressed as mean ± standard deviation (SD). Contracted, granular flattened and degranulated flattened represent the morphology classes of the amebocyte cells.

The multivariate model was significant for amoebocyte cell density (*F*=39.77, *p*<0.001), viability (*F*=196.61, *p*<0.001), contracted (*F*=2.327, *p*<0.05), granular flattened (*F*=7.438, *p*<0.001) and degranulated flattened (*F*=2.246, *p*<0.05). No significant difference (*F*=1.139, *p*>0.05) observed in cell density between captive group and wild group of *T. gigas* (mean value of 0.883×10^7^ cells/mL and 0.917×10^7^ cells/mL, respectively) (Figure 3). This indicated that the diet (*Meretrix meretrix*) and captive conditions in our study was suitable for *T. gigas* to reproduce amoebocyte cells, hence the blood cell count was restored.

The next parameter investigated was cell viability, which was aimed at assessing the quality of amoebocyte cells rather than their quantity. Regardless of the anticoagulant solution used, the results showed significant reduction in cell viability (*F*=808.075, p<0.001) of captive crabs compared to the wild crabs (Figure 3). Figure 2 shows amoebocyte cells’ density and viability of wild horseshoe crab under microscope.

**Figure 2.**
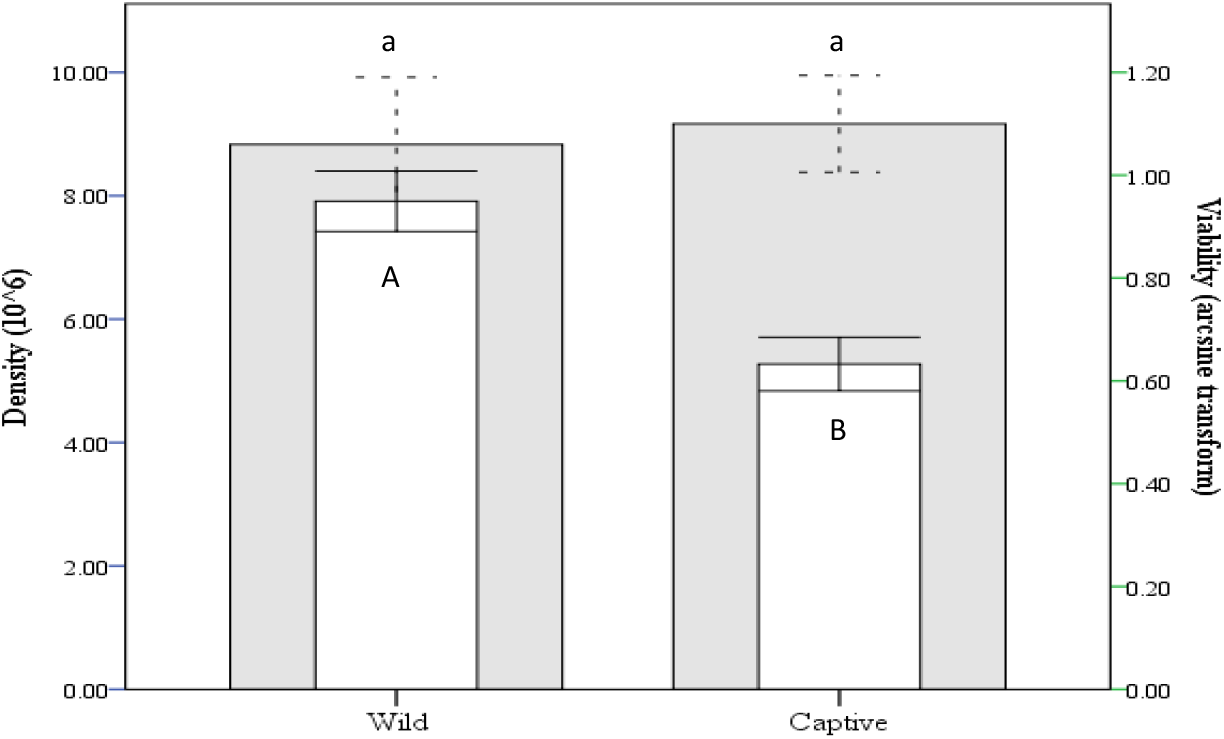
Comparison of amoebocyte cell density and viability between wild and captive *T. gigas*. Grey bars represent density and White bars represent viability. Standard error bars are shown as complete I-beam for density and dashed I-beam for viability. Post-hoc results are shown in lower case letters for density and upper case for viability.

**Figure 3.**
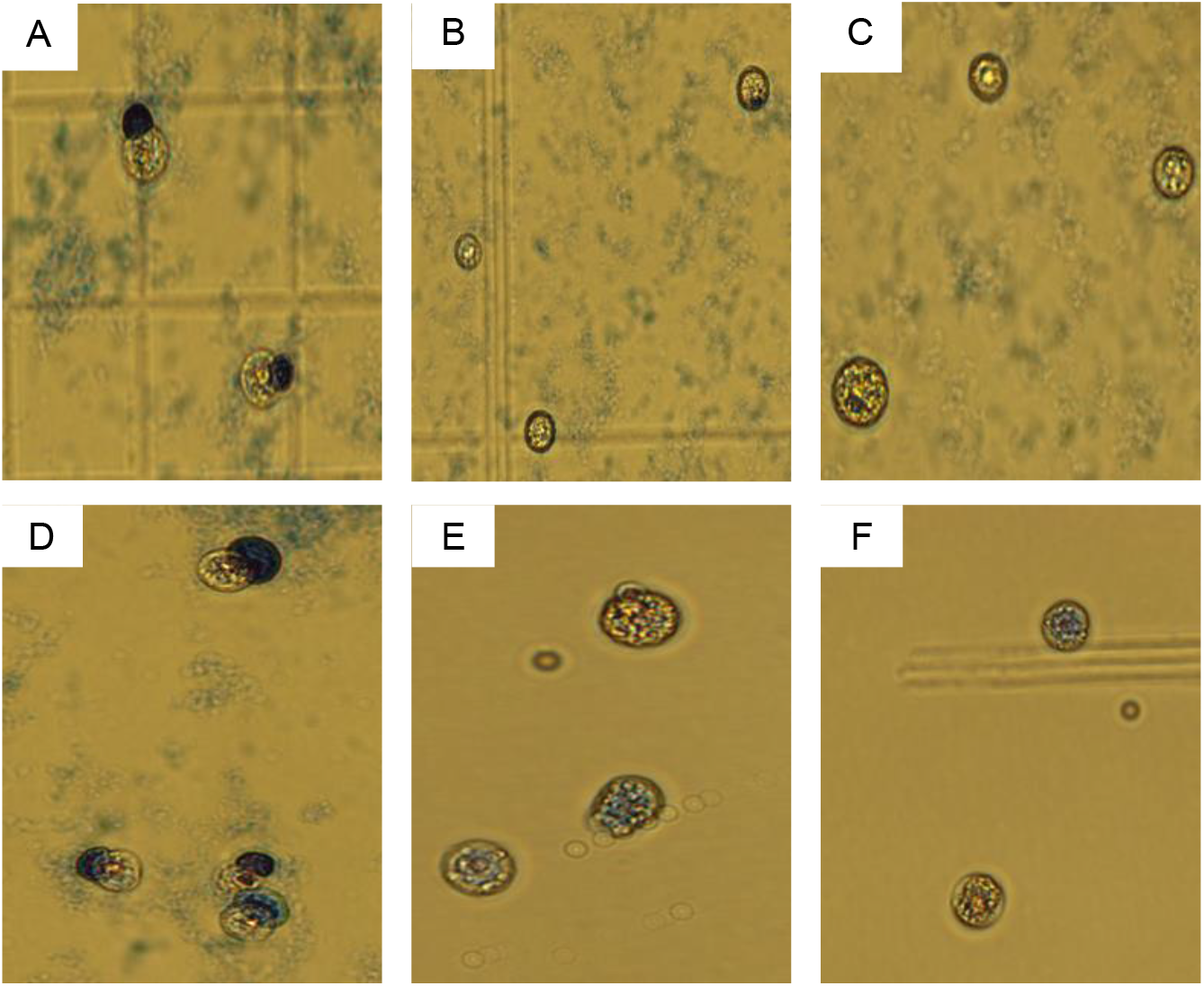
Amoebocyte cell viability assay of wild horseshoe crab. A to F: Anticoagulant formulas (40x magnification). Dead cells appear blue/dark cell, while live cells appear yellow/bright.

Decline in quality of hemolymph generally and amebocyte cells specifically, and even horseshoe crab mortality have been previously linked to captivity related issues. Infectious diseases such as blue-green cyanobacteria, fungi, Gram-negative bacteria and various types of parasites can cause health to decline (Nolan & Smith, 2009). Erosion of the carapace is also an added indicator of parasitic infections. In prolonged captive conditions, such as employed in this study, non-infectious diseases such as deficiencies in certain dietary elements (Carmichael & Brush, 2012), limited movement in culture tanks and absence of tidal rhythms also affects hemolymph quality (Kwan et al., 2014; Nolan & Smith, 2009).

This study recorded zero mortality during the study period, no observed infections or carapace erosion. Hence, the decline in amebocyte cells viability might be due to dietary deficiencies and other captivity related issues. Some studies have shown that diet has a significant impact on the gut endocrine cells in terms of regulating specific hormone release that eventually helps the animal to fight against invading pathogens and disease vectors (El-Salhy, Mazzawi, Hausken, & Hatlebakk, 2016; Webster, Dircksen, & Chung, 2000; Zudaire, Simpson, & Montuenga, 1998). Perhaps a more diverse and complete diet could maintain high quality amebocyte cells in captivity. The sampling site (Balok beach) is a nesting ground of both *T. gigas* and *Carcinoscorpius rotundicauda* (CR) and is rich in various benthic organisms such as mollusks, insects, crustaceans, and polychaetes. Previous studies had also shown that the gut content analysis of *C. rotundicauda* showed that besides bivalves, gastropods, crustaceans and polychaetes, CR also consumed other miscellaneous food items and even plant materials. The feeding of horseshoe crab has also been linked to the gender size, age, season, etc (Botton, 2009; John, Kamaruzzaman, Jalal, & Zaleha, 2012; Lewbart, 2011).

The anticoagulant formulations significantly affected all tested parameters namely amoebocyte cells’ density, viability, percentage of contracted cells, granular flattened and degranulated flattened cells (*p*<0.001). Significant interaction was observed between anticoagulant and captivity (*p*<0.001), hence, the effect of anticoagulant was re-analysed using data from wild crabs only in order to eliminate the effect of captivity.

Another parameter observed was the percent coefficient of variation (CV %) within the replicates. Amebocyte cell density varied by 18% in both the wild and captive. In the viability test, wild crabs varied by only 2%, while captive crabs varied by 23%. In morphology analysis, average variation was 9% in the wild group and 20% in the captive group. These results indicated that the 18 crabs showed different levels of adaptation to captivity.

### Effect of anticoagulant composition on horseshoe crab’s hemolymph quality

Besides the health status of the horseshoe crab, coagulation can also impair the cell viability, hence, anticoagulants are always used when collecting blood. Anticoagulants of choice are usually isotonic solutions and contain salts, xanthine derivatives (e.g. caffeine) or other agents (e.g. N-ethylmaleimide) that can modify sulfhydryl groups (-SH). An ideal anticoagulant act as a chelating and buffering agent with pH in the range of 4.5-8 (Morales et al., 2018; Rinkevich, 2012). The coagulation cascade in horseshoe crab hemolymph is dependent on pH and temperature. It is crucial to inhibit this enzymatic reaction triggered by a serine protease, mainly factor C (Ariki et al., 2004; Novitsky, 2009; Rinkevich, 2012; Young, Levin, & Prendergast, 1972). An ideal anticoagulant will show high viability, high percentage of granular cells (C and GF) and a low percentage of degranulated cells (DF).

The effect of different anticoagulants formulations on cell viability was apparent in this study. Amebocyte cell viability of wild crabs in formula A, B, C, D, E and F were 80%, 84%, 86%, 36%, 45% and 54%, respectively. Captive crabs had also shown the same trend with viability percentage of 40%, 52%, 58%, 11%, 18%, 39%, respectively. The anticoagulant formulations significantly affected all tested parameters namely amoebocyte cell density, viability, percentage of contracted cells, granular flattened and degranulated flattened cells (*p*<0.001). Significant interaction was observed between anticoagulant and captivity (*p*<0.001), hence, the effect of anticoagulant was reanalysed using data from wild crabs only in order to eliminate the effect of captivity. The new GLM was significant for amoebocyte cells’ density (*F*=59.962, *p*<0.001), viability (*F*=314.166, *p*<0.001), contracted (*F*=268.79, *p*<0.001), granular flattened (*F*=3772.932, *p*<0.001) and degranulated flattened (*F*=314.166, *p*<0.001). The anticoagulant formulations still significantly affected all tested parameters. In multiple pairwise comparisons using Tukey post-hoc test, Anticoagulant B and C showed significantly higher viability in both captive and wild horseshoe crabs (p<0.001), however, no significant difference between Formula B and C (p=0.395) (Figure 4, 5 and 6).

**Figure 4.**
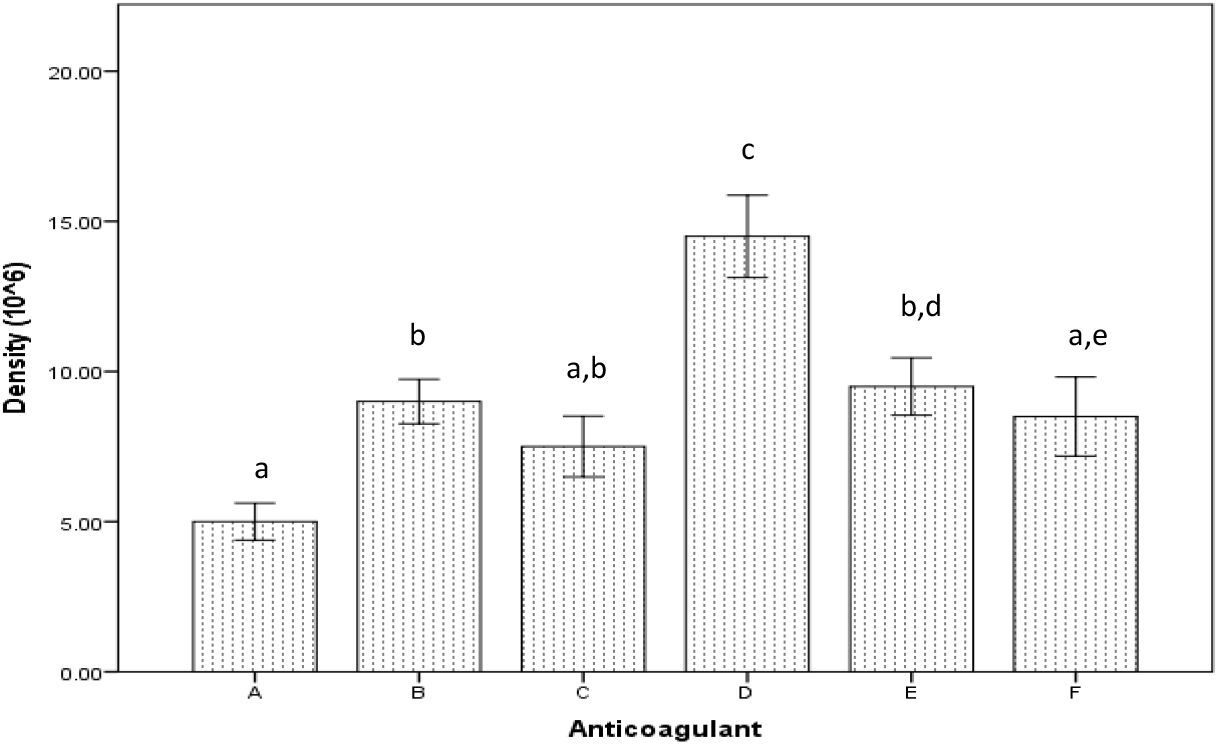
Effect of anticoagulant formulations on amoebocyte cells’ density in wild *T. gigas*. Standard error bars are shown and post-hoc results are shown in different alphabets.

**Figure 5.**
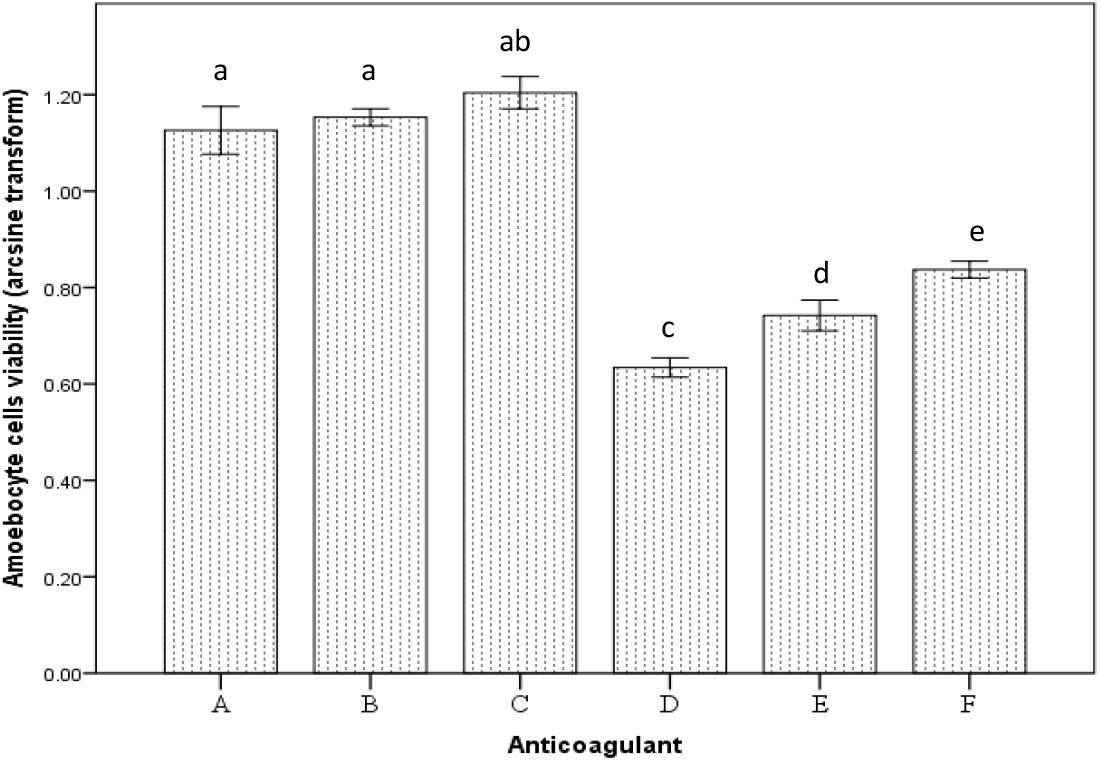
Effect of anticoagulant formulations on amoebocyte cells’ viability in wild *T. gigas*. Standard error bars are shown and post-hoc results are shown in different alphabets.

**Figure 6.**
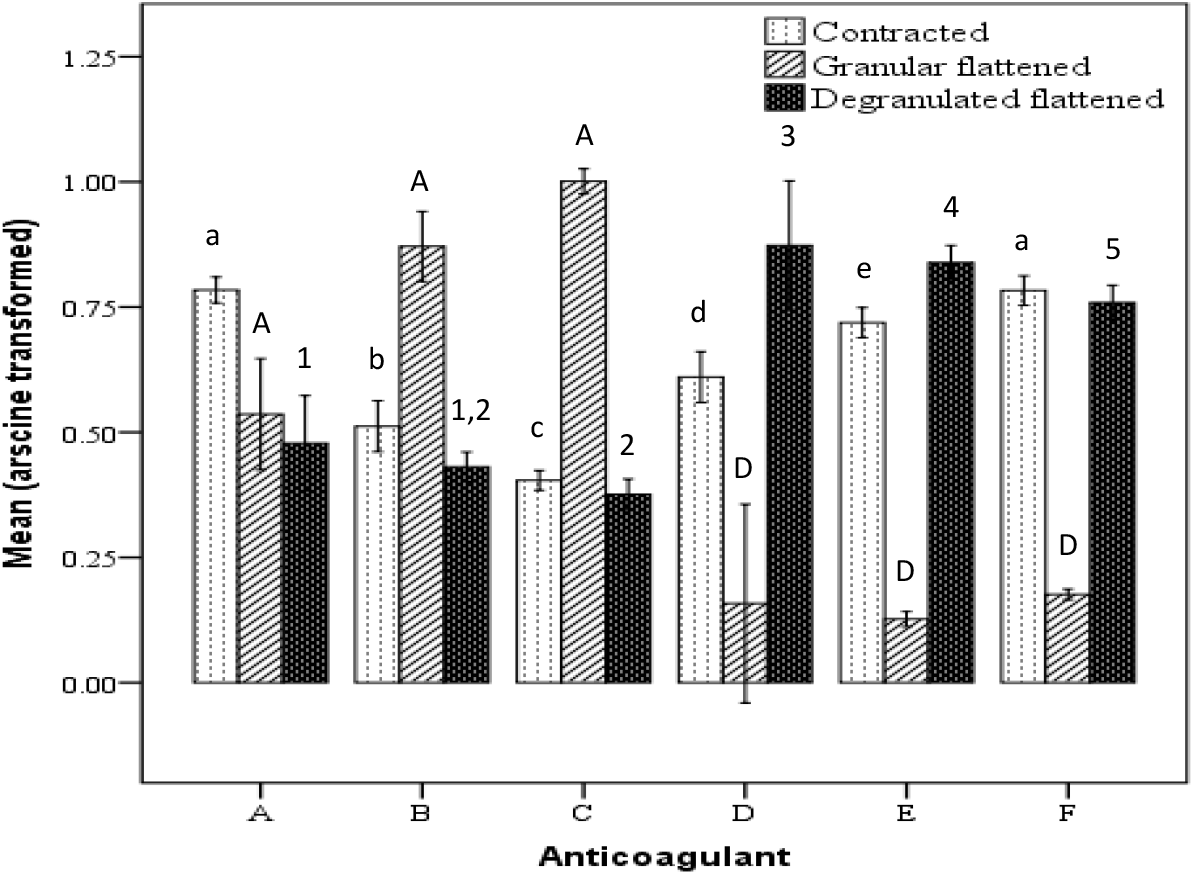
Effect of anticoagulant formulations on percentage of contracted (C), granular flattened (GF) and degranulated flattened (DF) cells in wild *T. gigas*. Standard error bars are shown and post-hoc results are shown in lower case letters for C cells, upper case letters for GF cells and numbers for DF cells.

The new GLM of only wild group data was significant for amoebocyte cell density (*F*=59.962, *p*<0.001), viability (*F*=251.388, *p*<0.001), contracted (*F*=71.949, *p*<0.001), granular flattened (*F*=62.469, *p*<0.001) and degranulated flattened cells (*F*=39.67, *p*<0.001). In multiple pairwise comparisons using Tukey post-hoc test, density analysis showed that A, B, C and E were similar, indicating similar chelation capacity. However, in the viability analysis, A, B and C were significantly better than all other formulas. Formula C was also significantly better than A in a pairwise comparison. Formula C showed significantly higher cell density that all other formulas due to cell clumping (*p*<0.001) (Figure 4). Cell viability analysis showed that formula A, B and C were significantly higher than in other formulas. Formula C was significantly higher than formula A in a pairwise comparison (*p*<0.01). No significance difference in viability was observed between formula B and C (*p*>0.05) (Figure 5). In morphology analysis, formula B and C showed significantly lower percentage of C cells (*p*<0.001) and significantly higher GF (*p*<0.001) compared to others, however, both cell types are considered to be functional. DF cells are considered dead and non-functional was significantly high in formula D, E and F. This showed that formula A, B and C were able to maintain amebocyte cell functionality. No significant difference in percentage of DF cells between formula A, B and C (*p*>0.05). Overall results showed that formula B and C are better than other formulas, but formula C is slightly better than B (Figure 6). The effect of different anticoagulants formulations on cell morphology was similar to cell viability. The performance of different anticoagulants used in this study can be summarized as B>C>A>F>E>D.

All top three anticoagulants formulations (A, B and C) were adjusted to pH 4.6 that inhibits serine protease zymogen which in turn initiates coagulation and were all supplemented with chelation agent and energy source. The isoelectric point of serine protease is 4.6. The selected 4.6 pH also deactivates endotoxins which help in further coagulation inhibition (Ribeiro et al., 2010; Soares et al., 2012). These three formulas differed in buffering capacity and osmotic pressure regulation. Formula B and C contained citric acid which can serve as both chelation and buffering agent, while formula A did not contain citric acid or any buffer. Formula B had NaCl and Citric Acid, while formula C had Sodium Citrate and Citric Acid. Therefore, formula C had the best buffering system as it contained a good divalent cation which are good buffering system (Armstrong & Conrad, 2008; Lee & Arepally, 2012). While NaCl in formula A and B is good in maintaining osmotic pressure (Ratcliffe, 1993), buffering capacity appeared to be more important. Formula B compensated for the lower buffering capacity with higher citric acid content which might have enhanced the chelation, while NaCl maintained the osmotic pressure. Therefore, simplicity of formula A and its lack of chelation and buffering capabilities might have led to lowest cell viability percentage among tested formulations. These results were in agreement with previous study on *Penaeus japonicus* where low pH citrate–EDTA anticoagulant buffer prevented coagulation where EDTA inhibited the prophenoloxidase (proPO) activation and citric acid delayed cell breakdown which ultimately maintained cell integrity and cell viability (Johansson, Keyser, Sritunyalucksana, & Söderhäll, 2000).

The last anticoagulant formula that was adjusted to pH 4.6 was formula D. The formula was solely based on a study by Liang and Liu (1982) who reported that proclotting enzymes can be inhibited by dimethyl sulfoxide (DMSO). However, this anticoagulant formula exhibited the poorest performance in both wild and captive horseshoe crab. This could be due to the absence of a buffering system and limited chelation agent of 1 mM Na_2_EDTA compared to 9 −10 mM in other formulas.

Anticoagulant E and F were studied based on previous reports on amebocyte cell culture which indicated that amebocyte cells appeared at neutral pH (7.2-7.8) in gill flaps cultures where they are produced (Armstrong, 1979; Friberg, Weathers, & Gibson, 1992; Gibson & Hilly, 1992; Joshi, Chatterji, & Bhonde, 2002). Therefore, anticoagulant E and F were set to fall within optimum pH (6-9) for serine protease activity (Cuervo et al., 2008). The performance of these 2 anticoagulants was to verify whether it is more important to inhibit the serine protease or to mimic the natural chemical environment of amebocyte cells and provide good buffering systems. Formula E consisted of a universal Good’s buffer (HEPES), while formula F was made up of Bicarbonate-Carbonate which is also a common and efficient buffer. Both buffers were also set to their optimum pH level in which their buffering capacity is at peak performance. The results had shown that it is more important to deactivate serine protease zymogen.

Recent attempt by Tinker et al., (2020) to use aquaculture horseshoe crabs for LAL production has increased number of concerns. For instance, the anticoagulant formulation adopted in their study consisted of a mixture of N-ethylmaleimide (0.125%), NaCl (3%), and Tris-HCl (0.5 M, pH 7.5) extracted from the previous works (Armstrong & Conrad, 2008; Coates, Bradford, et al., 2012; Coates, Whalley, et al., 2012) for amebocyte density analysis failed to address the key information such as pH of the anticoagulant mixture and the accuracy of the novel anticoagulant constituents adopted from the earlier work (Armstrong & Conrad, 2008). We assume that further modification and optimization on formula B and C used in this study could improve the cell viability. Generally, the results proved that captivity had a profound negative effect on cell viability and morphology. The results also indicated the importance of adequate energy supply to the cells in order to maintain cellular activities, as well as buffering system that can maintain cell membrane integrity.

## CONCLUSION

To conclude, captive breeding has a profound negative impact on the amebocyte viability, therefore, captive rearing conditions must be improved in terms of developing nutritionally enriched feed for horseshoe crab to achieve better amebocyte quality. Anticoagulant formulation had profound effects on amebocyte cell density, viability and morphology. In terms of anticoagulant agents, formula B and C showed good anticoagulant activity and could maintain the viability of amebocyte cells. Overall, hemolymph quality can be used as an indicator to determine the rearing condition and health status of horseshoe crabs under prolonged captivity.

## ACKNOWLEDGEMENT

Special thanks to Prof. Mark L Botton for his technical advice and manuscript editing. Authors extend their sincere thanks to two anonymous reviewers for their expert advice to improve the manuscript. This project was funded by Malaysian government under the Fundamental Research Grant Scheme (FRGS 15-210-0451 and FRGS 19-.021-0629).

## AUTHORS CONTRIBUTION

AJ: Idea conceptualization, data analysis, Manuscript preparation

HS: Manuscript preparation and laboratory work, data analysis and interpretation

SI: manuscript review, experimental design

KY: Formal analysis and investigation

---

**Supplementary File 1:** Morphometric measures of experimental crabs used in the captivity study.

| **Sex** | **Total Length (cm)** | **Carapace Width (cm)** | **Interocular Width (cm)** | **Telson Length (cm)** | **Weight (g)** |
| --- | --- | --- | --- | --- | --- |
| **Female** | 38.8 | 21.4 | 11.1 | 17.5 | 575 |
| **Female** | 38.7 | 21.7 | 10.8 | 17.1 | 655 |
| **Female** | 43.5 | 23.1 | 12.4 | 19.2 | 730 |
| **Female** | 40.6 | 21.4 | 11.6 | 17.7 | 665 |
| **Female** | 42.7 | 23.3 | 12.4 | 18.7 | 860 |
| **Female** | 42.3 | 23.5 | 12.2 | 18.6 | 820 |
| **Female** | 43.4 | 23.1 | 12.7 | 19.1 | 810 |
| **Female** | 47.7 | 22.4 | 12.6 | 23.8 | 980 |
| **Female** | 45.5 | 22 | 12 | 25 | 880 |
| **Female** | 42.5 | 21.3 | 11.4 | 23.1 | 820 |
| **Female** | 37.6 | 20.5 | 9.9 | 16.4 | 625 |
| **Female** | 45.4 | 22 | 12.2 | 22.4 | 955 |
| **Female** | 41.6 | 20.5 | 11.3 | 21.7 | 845 |
| **Female** | 42.3 | 19.7 | 12.3 | 19.7 | 705 |
| **Female** | 41.8 | 21.7 | 10.3 | 17.9 | 675 |
| **Female** | 46.4 | 21.2 | 10.6 | 22.1 | 705 |
| **Female** | 41.2 | 21.4 | 11.8 | 17.9 | 785 |
| **Female** | 49.2 | 24.5 | 14.2 | 24.5 | 920 |

